# A short remark on Ontogenetic model and its analytical solution

**DOI:** 10.1101/031732

**Authors:** Ya Li, Ji-Huan He

## Abstract

The ontogenetic growth modeling has been of conspicuously scientific interest, as well as its practical importance in many industrial activities for optimal design. Much work has been done recently to understand the growth mechanism of living organism with a nonlinear time dependence. Some quantitative metabolic models which have been provided to describe organism growth curves are in comparison with what for ontogenetic growth represented by G.B. West et al. (West et al., 2001) in his basic formulations used to discuss the application on different growth patterns. Herein the analytical solution is also involved. In this paper we re-analyzed the assumptions underlying them to find whether the basic relation of organism growth estimated can inosculate with the equation we have proposed.

## Introduction

As the individual’s body size increases, its mass – relative to the time changes. There are so many biological processes from cellular metabolism to organism dynamics and the relationships between size and rate are characterized by particular growth rhythm. More physiological approaches to describe life growth curves have drawn on more sophisticated models for ontogenetic growth.

The von Bertalanffy equation used to describe the ontogenetic growth of organisms until West et al. (West et al., 2001) provided a model to describe the ontogenetic growth. The dependence of a biological variable B on body mass M is typically characterized by nonlinear system of equations. The base metabolic rate B of numerous species (both warm blooded and cold-blooded organisms) is empirically observed to be proportional to body mass m or volume with an exponent of 0.75, i.e. B = B_0_m3/4 (West et al., 2001), where B_0_ is a constant that is characteristic of the type of organism. Similarly, the advantages of the logistic model over the ontogenetic growth are the wide adaptability for many species including animal and plant growth (Shi et al., 2013). Obviously, many contending models emerged describing the relationship between mass and time (Shi et al., 2013). Among these models, the von Bertalanffy equation might be the most popular one (Bertalanffy, 1957). Traditionally, these variables have been related to a single, easily measurable biological attribute: body size (Calder, 1984). A quantitative model has also been developed to describe ontogenetic growth, taking into account a single average cell in the building block, then the rate equations for a, b and m concentrations can be written as:

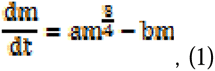

where the constants a and b can be related to physical properties of the single average cell, m represents the body mass at time t. It claimed that 0.75 exponent had been well supported by the data on many biological spcies like mammals (Rogers et al., 1995), birds (Weathers and Siegel, 1995), fish (Xie and Sun, 1990; Brett, 1989), mollusks (Hamburger, et al., 1983), chicken (Nagai, et al., 2015) and plants (Enquist, et al., 1998). Then P.J. Shi et.al. tested the generality of the ontogenetic growth on different crop species and compared the goodness-of-fits between the ontogenetic growth and other non-nonlinear models that are often used to describe the ontogenetic growth (Shi et al., 2013). However, they noticed some ignorance of crops existed. In this work we have derived a theoretical framework for the growing trajectory in the context to elucidate the growth of biological species.

## Materials and methods

### Data

The body mass data of several spcies were extracted from the research of the general model by West et al. and m_0_ is the intial mass at birth (t=0), M represents the asymptotic maximum body mass, which is ultimate constant. They can be written as:

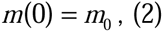

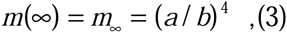

While these metabolic models provide a valuable insight into ontogenetic growth, further refinements must capture the relevant relation in the architecture of biological species. Ultimately, ontogenetic growth models must also include terms to describe the increase in mass associated with time.

Values of a, m_0_ and M for these and several other species had been provided during these research. They mostly explained the sigmoidal time dependence of the mass of the biomass (Ulmschneider and Searson, 2015). Analysis of ontogenetic growth and in vivo data of animal models and humans had been used to suggest that the growth follows a universal growth law with three adjustable parameters: the initial and final mass of the living beings, and the constant *a,* whereas b (=a/M^1/4^) should scale as aM^-1/4^. The current challenge in this field is to bring physical insight into this universal behavior by combining models (West, 1997).

### Models

Consistent with our predictions, we will verify the equations to see whether we gave which had been arranged as Eq. (7) is valid when compared to match if it can fit the biological species. Here, what all the parameters represents have been referred above. Consequently, the derivative of the functions can be got step by step as the following formulas. To differentiate the implicit function of Eq. (1) as

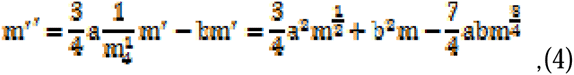

with intial conditions of Eq. (1) and Eq. (4):

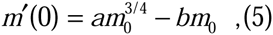

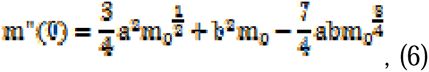

We propose a new hypothetical ontogenetic growth model to express the law of mass – relative to the time, then we have

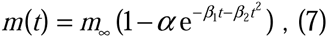

with intial conditions of Eq. (7):

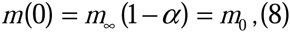

Differentiating Eq. (8) with respect to time results in

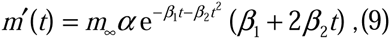

Similarly we have the following two-order differential equation by differentiating Eq. (9) again:

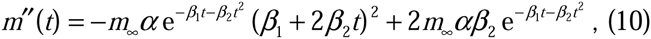

Integration of Eq. (5) and Eq. (9) with initial information when t=0 results in

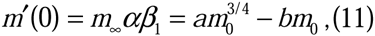

By incorporating Eq. (6) into Eq. (10) with initial information when t=0, we have the following relationship:

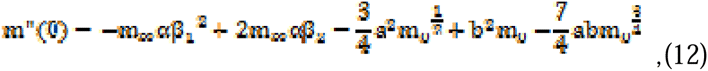

Meanwhile we have the known formula got by Eq. (8) as

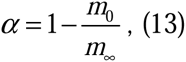

After the above steps, we can get the parameter:

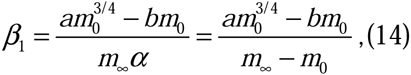

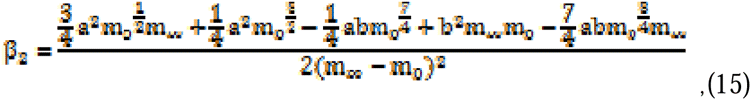

### Relationship between two models

Briefly, in this work we consider classical methods of ontogenetic growth together with mixtures theory with a rigid individual. The examples in general equations, were gave as the figures as below. Besides, The data was chosen from the samples of general equations by West et al. (West et al., 2001). In order to prove the relation with mass and time, we have set out to determine whether the equation is suitable for the nonlinear dependence.

Put the data in he equations we gave, then we found that all can fit the data extremely well (Fig. 1, Fig. 2, Fig. 3 and Fig. 4) by using Eq. (7).

(1) Guinea pig: *m*_0_=5, M=840, a=0.2, b=0.04, α=0.99, β_1_=0.000578, β_2_=0.0000167

(2) Guppy: *m*_0_=0.008, M=0.15, a=0.1, b=0.167, α=0.947, β_1_=0.01, β_2_=0.00045

(3) Hen: *m*_0_=43, M=2050, a=0.502, b=0.075, α=0.979, β_1_=0.0026, β_2_=0.0000366

(4) Cow, *m*_0_=33333, M=442000, a=0.276, b=0.01, α=0.925, β_1_=0.00085, β_2_=0.00000262

**Fig. 1.**
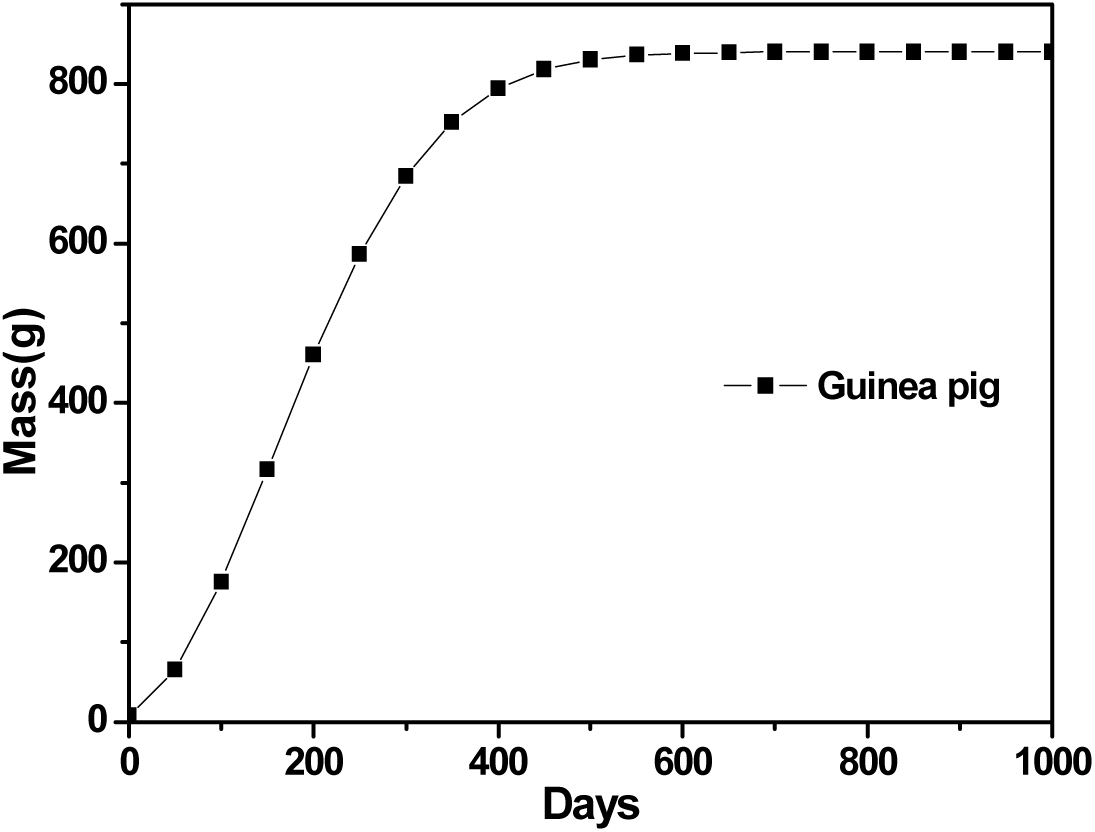
Guinea pig typical example of fits to growth curve (solid lines) using Eq. (7).

**Fig. 2.**
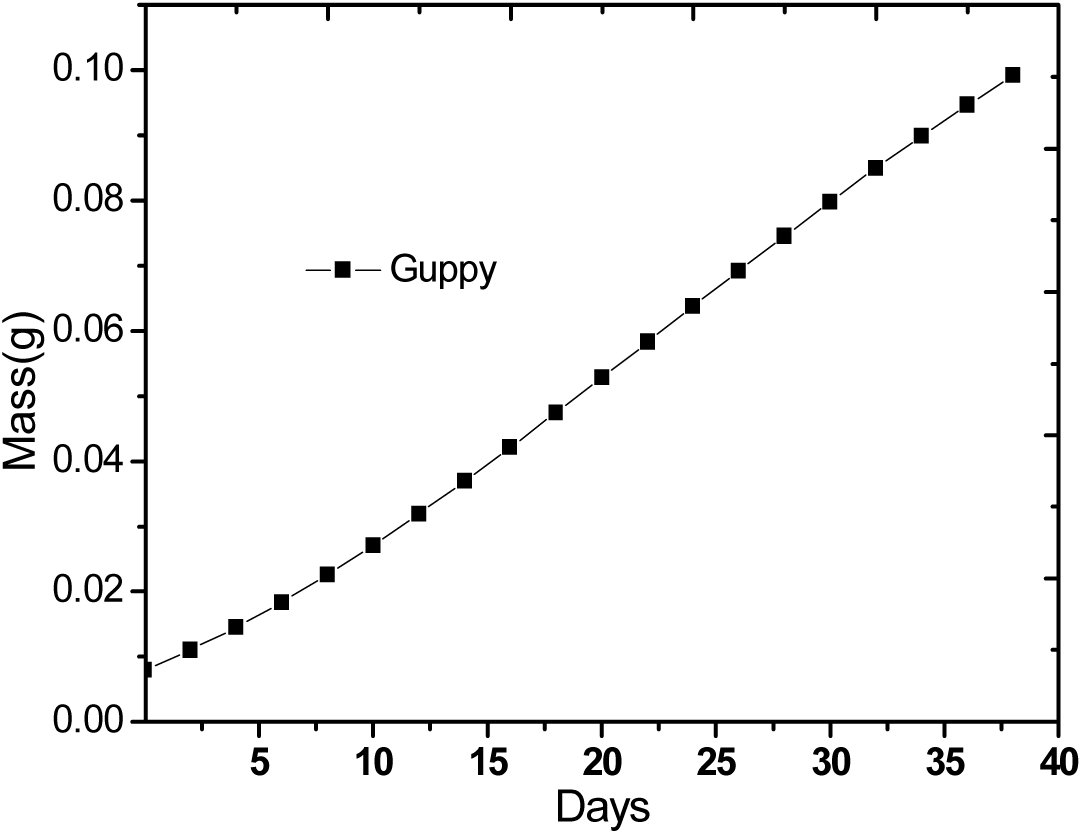
Guppy typical example of fits to growth curve (solid lines) using Eq. (7).

**Fig. 3.**
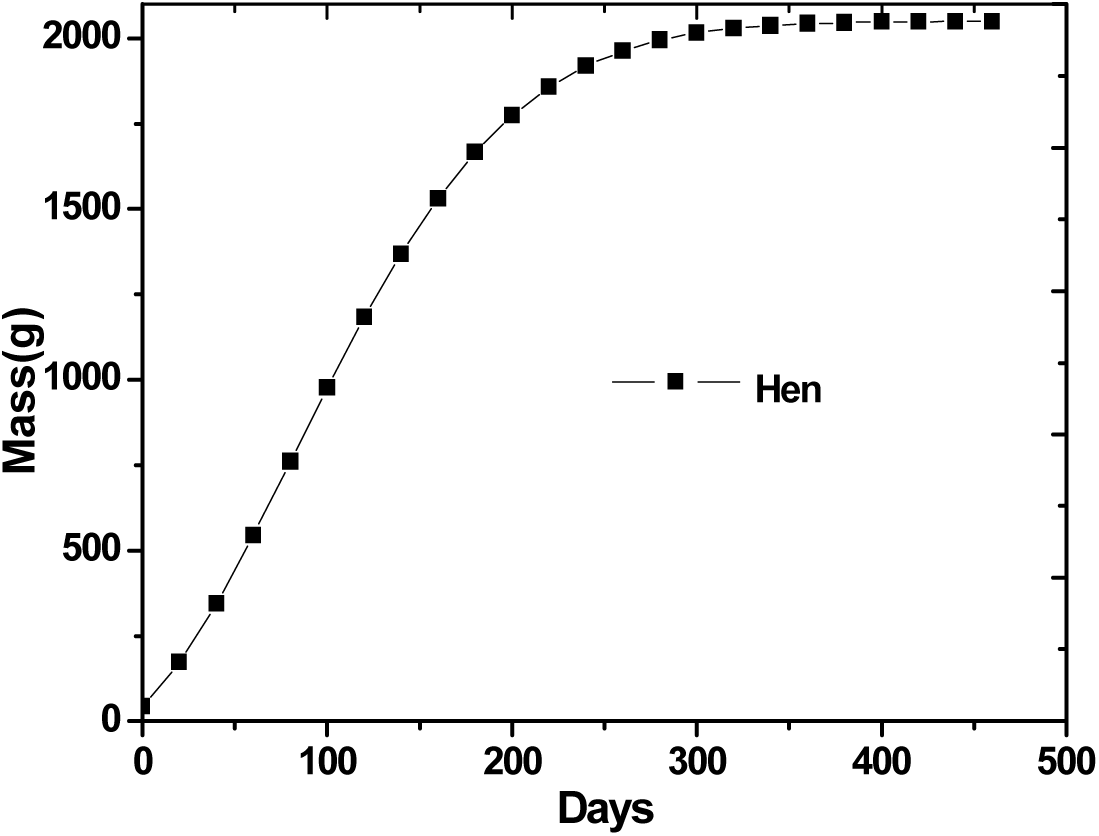
Hen example of fits to growth curve (solid lines) using Eq. (7).

**Fig. 4.**
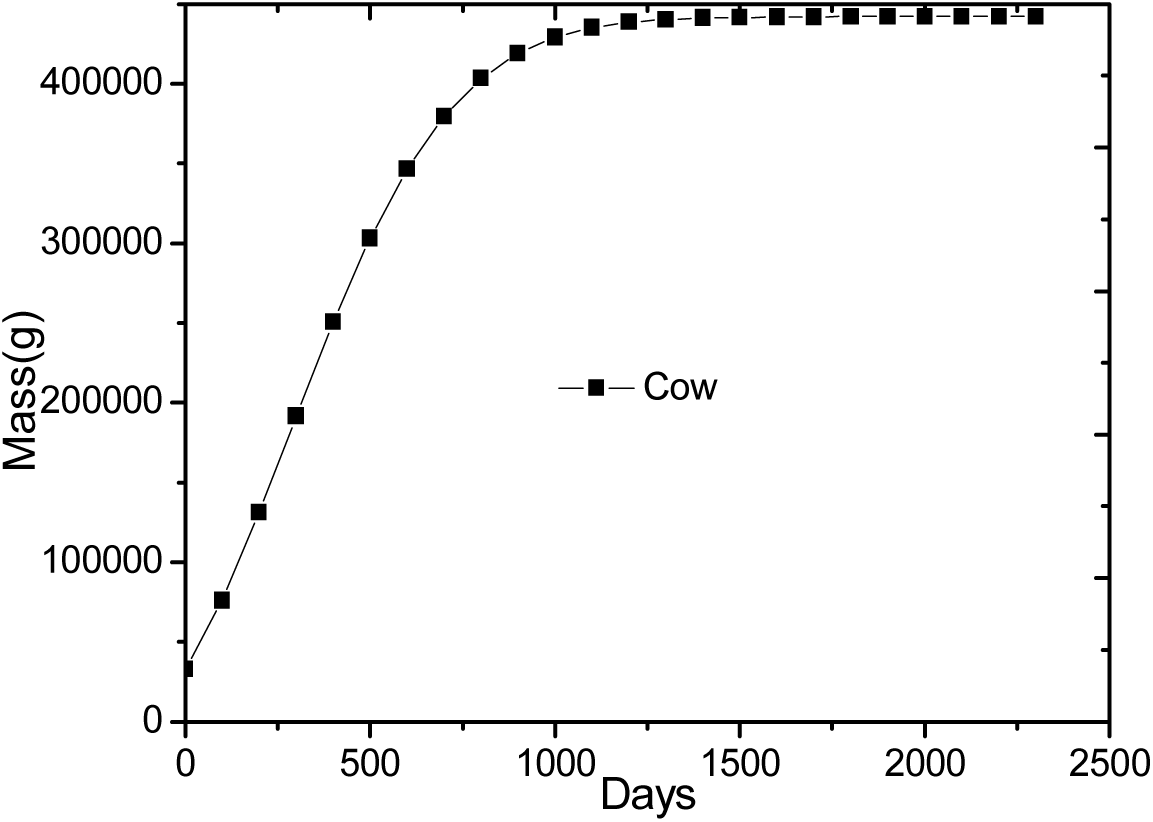
Cow typical example of fits to growth curve (solid lines) using Eq. (7).

## Conclusion

In summary, the equation of the mode we gave meets all the samples used in the so-called “general” ontogenetic growth model and they all satisfy the nonlinear dependence. The model structure is simple and typical, it has wide adaptability for many species including animals and crops growth. We get the same shape of the general growth equation from nature.

## Acknowledgments

The work is supported by Priority Academic Program Development of Jiangsu Higher Education Institutions (PAPD), National Natural Science Foundation of China under grant No.11372205 and Project for Six Kinds of Top Talents in Jiangsu Province under grant No. ZBZZ-035, Science & Technology Pillar Program of Jiangsu Province under grant No. BE2013072.

